# *À la carte*, *Streptococcus pneumoniae* capsular typing: using MALDI-TOF mass spectrometry and machine learning algorithms as complementary tools for the determination of PCV13 serotypes and the most prevalent NON PCV13 serotypes according to Argentina’s epidemiology

**DOI:** 10.1101/2022.12.22.521467

**Authors:** Jonathan Zintgraff, Florencia Rocca, Nahuel Sánchez Eluchans, Lucía Irazu, Maria Luisa Moscoloni, Claudia Lara, Mauricio Santos

**Author notes:** (Florencia Rocca), (Nahuel Sanchez Eluchans), (Maria Luisa Moscoloni), (Claudia Lara), (Mauricio Santos).

## Abstract

Laboratory surveillance of *Streptococcus pneumoniae* serotypes is crucial for the successful implementation of vaccines to prevent invasive pneumococcal diseases. The reference method of serotyping is the Quellung reaction, which is labor-intensive and expensive.

In the last few years, the introduction of MALDI-TOF MS into the microbiology laboratory has been revolutionary. In brief, this new technology compares protein profiles by generating spectra based on the mass to charge ratio (m/z).

We evaluated the performance of MALDI-TOF MS for typing serotypes of *S. pneumoniae* isolates included in the PCV13 vaccine using a machine learning approach. We challenged our classification algorithms in “real time” with a total of new 100 isolates of *S. pneumoniae* from Argentinian nationwide surveillance.

Our best approach could correctly identify the isolates with a sensitivity of 58.33 % ([95%IC 40.7-71.7]); specificity of 81.48 % ([95%IC 53.6-79.7]); accuracy of 63.0% ([95%IC 61.9-93.7]); PPV of 80.77% ([95%IC 64.5-90.6]) and NPV of 59.46% ([95%IC 48.9-69.2]).

In this work, it was possible to demonstrate that the combination of MALDI-TOF mass spectrometry and multivariate analysis allows the development of new strategies for the identification and characterization of Spn isolates of clinical importance; and we consider that by using AI, as more data becomes available the models will get better and more precise.

## Introduction

*Streptococcus pneumoniae* (Spn) is a human pathogenic microorganism **(1)**, responsible for a wide spectrum of infectious diseases and invasive processes that constitute an important cause of morbidity and mortality in the world. Infectious diseases occur when the bacteria gain access to generally sterile areas of the respiratory tract, thus producing the dissemination, colonization and invasion of pneumococcus, mainly the middle ear (producing otitis media), the lungs (pneumonia), the bloodstream (bacteremia) or the central nervous system (meningitis). The bacteremia that is related to higher mortality and morbidity is generally caused by complications of pneumonia. Spread between different individuals occurs by direct contact with secretions from colonized individuals **(2)**.

According to estimates by the World Health Organization (WHO), sepsis, neonatal and community pneumonia accounted for 15% of annual deaths in children under 5 years of age in 2015, with *S. pneumoniae* being the most common cause of pneumonia in both developed and developing countries **(3–5)**. The incidence of invasive pneumococcal disease (IPD) varies geographically, and is highly related to poverty **(3)**.

The antigenic diversity of the capsular polysaccharide allows pneumococcus to be classified into different serotypes. The reference technique for serotyping was described by Neufeld in 1902 and is called the Quellung Reaction **(6–8)**. Briefly, this technique utilizes a microscope and specific pneumococcal antisera and is commonly used in reference and research laboratories worldwide. This method uses a chessboard system, in which the pneumococcus is sequentially tested with antisera pools until a positive reaction is observed. Each pool contains different mixtures of antisera against pneumococcal serotypes. Once a pool is established, the individual type and group antisera that are included in the reactive pool are tested individually in sequence in order to determinate the final serotype.

Contemporary studies led to the development of a system that allowed serogroups to be distinguished from serotypes:

a. Serotype: it is defined as an isolate that produces a polysaccharide with unique chemical and immunological properties.
b. Serogroup: it is defined as a group of serotypes that share several immunological properties or antigenic determinants (cross-reactions).

Currently, 100 immunologically different capsular serotypes have been described **(9)**. The importance in determining the pneumococcal capsular serotype is fundamentally based on the fact that its distribution is related to a series of factors, such as age, sex, clinical picture, geographical area, antibiotic sensitivity and many other epidemiological data that allow correlate with the prevalence of certain serotypes. On the other hand, given that it is a vaccine-preventable disease and that the vaccines developed and implemented are directed at specific capsular serotypes, the determination of circulating serotypes is extremely important.

Pneumococcal vaccines provide serotype-specific protection, it is important that vaccines prevent disease caused by the most clinically relevant serotypes. Therefore, vaccines provide the greatest impetus for recognizing capsular diversity and serotype epidemiology. Pneumococcal vaccines are an important public health tool and have undergone dramatic changes in recent years. Therefore, there are many excellent overviews of pneumococcal vaccines **(10–12)**.

In Argentina, in 2012, the pneumococcal conjugate vaccine, PCV13 (which includes purified capsular polysaccharide for serotypes 1, 3, 4, 5, 6A, 6B, 7F, 9V, 14, 19A, 19F, 18C, and 23F) was included in the National Immunization Program in a 2 + 1 schedule (2, 4, 12 months) **(13)**.

The impact of PCV13 introduction on serotype distribution may vary with time and location, as the epidemiology of serotypes is quite variable both geographically and temporally. In general, conjugate vaccines have been effective in preventing IPD and carriage while offering variable immunity against cross-reactive serotypes. Nevertheless, several studies have shown that the introduction of PCVs has resulted in changes in epidemiology of pneumococcal diseases and can drive an increase in the frequency of preexisting resistant variants of non-vaccine serotypes due to the removal of competition from vaccine serotypes **(14–20)**.

MALDI-TOF/MS (MS) is a new technology that in recent years, has acquired great importance in the identification of pathogens that are clinically relevant in public health **(21–23)**. The technique is based on the analysis of protein spectra containing peaks with an exactly determinable mass-charge ratio (m/z) generated by the impact of a laser on a previously crystallized isolate with an organic matrix. The potential of this methodology combined with machine learning algorithms for the detection of profiles in a wide variety of samples and its use as a screening technique is expanding, due to its low-cost and high performance **(24)**.

Although the initial investment in the purchase of a mass spectrometer is relatively high, the cost for the identification of a single isolate is lower than previously used biochemical or molecular techniques **(25,26)**. Among the advantages, the simplified workflow is one of the key points for the rapid acceptance of MS in laboratories and by the broad range of applications that it could be extended.

In this work, we proposed to explore the potential of MALDI-TOF technology as a complementary and screening tool for Spn capsular typing by developing classification models based on MALDI-TOF spectrometry in order to discriminate Spn strains from PCV13 and NON PCV13 isolates.

To achieve these two objectives, we created a spectral database with isolates of serotypes included in the PCV13 vaccine and, according to local epidemiology, the first 10 most prevalent NON PCV13 serotypes; we implemented unsupervised models from MALDI-TOF spectra to establish the basis for designing predictive classification models. Then we developed supervised classification models based on MALDI-TOF spectra that allowed to predict Spn isolates depending on whether they belong to the PCV13 or NON PCV13 class and finally we validated the models developed in the previous point with an independent set of samples.

These approaches will contribute to keep exploring the use of artificial intelligence in combine with traditional and nontraditional microbiological methods, in order to improve clinical diagnosis.

## Methods and Materials

### Bacterial isolates

A total of 23 isolates were selected, of which 13 corresponded to the so-called PCV13 (serotypes included in the PCV13 vaccine) and 10 non-PCV13 isolates (the most prevalent serotypes not included in the PCV13 vaccine). All of them belonging to the strain collection at the National Reference Laboratory for Meningitis and Bacterial Acute Respiratory Infections (NRL).

Prior to freezing, all isolates were identified using the optochin sensitivity and bile solubility tests. Pneumococci were serotyped using the Quellung reaction with sera provided by the Statens Serum Institute (Copenhagen, Denmark); isolates were stored at −70°C in Trypticase Soya (TS) broth with 15%–20% glycerol.

In this way, the 23 selected strains constituting the called “training set”. The information on the samples used can be seen in Table 1.

**Table 4.** Training set: isolates used for the creation of the machine learning models.

| Serotype | ID | Year | PCV13/NON PCV13 |
| --- | --- | --- | --- |
| 1 | 20380 | 2016 | PCV13 |
| 3 | 20117 | 2016 | PCV13 |
| 4 | 19202 | 2015 | PCV13 |
| 5 | 21128 | 2017 | PCV13 |
| **6A** | 20342 | 2016 | PCV13 |
| **6B** | 20372 | 2016 | PCV13 |
| **7F** | 23112 | 2019 | PCV13 |
| **9V** | 21329 | 2017 | PCV13 |
| **14** | 20437 | 2016 | PCV13 |
| **18C** | 23100 | 2019 | PCV13 |
| **19A** | 23049 | 2019 | PCV13 |
| **19F** | 21287 | 2017 | PCV13 |
| **23F** | 20018 | 2016 | PCV13 |
| **7C** | 23157 | 2019 | NON PCV13 |
| **8** | 23201 | 2019 | NON PCV13 |
| **11A** | 22167 | 2018 | NON PCV13 |
| **12F** | 23165 | 2019 | NON PCV13 |
| **13** | 23210 | 2019 | NON PCV13 |
| **15B** | 23148 | 2019 | NON PCV13 |
| **16F** | 23115 | 2019 | NON PCV13 |
| **22F** | 23143 | 2019 | NON PCV13 |
| **23B** | 23189 | 2019 | NON PCV13 |
| **24F** | 23207 | 2019 | NON PCV13 |

### Culture conditions

Starting from the isolates preserved at −70°C, sheep blood agar plates (5%) were inoculated with a small aliquot of them. After 24-48 hours, replication was carried out until growth of the order of 10^4^ CFU was obtained.

### Sample preparation and MALDI-TOF/MS spectra acquisition

Mass spectra acquisition was performed using a Microflex LT mass spectrometer (Bruker Daltonics, Germany) and the Flex Control software (version 3.4) with default parameter settings. Evaluation of the mass spectra was carried out using the Flex Analysis v3.4 software (BrukerDaltonics, Bremen, Germany) and MALDI Biotyper 3.1 software and library (version 9.0, Bruker Daltonics Germany).

All isolates were processed in triplicate in the same way and by the same operator, to minimize the variability of the methodology.

Seeding was performed by the direct method routinely used in clinical laboratories as indicated by the manufacturer **(27,28).**

One colony of the sample was placed in each well of the steel plate (MSP 96; BrukerDaltonics), then allowed to dry for a few minutes at room temperature and covered with 1 ul of the HCCA matrix (α-cyano-acid solution). 4-hydroxycinnamic acid diluted in 500 μL of acetonitrile, 250 μL of 10% trifluoroacetic acid and 250 μL of HPLC grade water). After dehydration of the sample-matrix mixture, the plate was introduced into the MicroFlex LT instrument and vacuum conditions were generated.

Mass spectra were recorded in the spectral region from 2000 to 20,000 Da (in linear positive ionization mode). The operation of the equipment was controlled by Flex Control v3.4 software. Each spectrum was a sum of 240 laser shots collected in increments of 40. Their recording was carried out at 40% of the maximum laser energy. The platform was pre-calibrated according to the manufacturer’s instructions using the BrukerDaltonics bacterial test standard (BrukerDaltonics, Bremen, Germany). The quality of the recorded spectra was evaluated using Flex Analysis software. All recorded spectra were stored in files (mzXML) for pre-processing and further analysis.

### Data analysis

All isolates were identified using the MALDI Biotyper RTC software and compared to the Bruker Biotyper database (reference library version 9.0). According to the manufacturer’s recommendations, the identification was considered reliable at species level when the score value was greater than 2.0 and at the genus level when the score value was between 1.7 and 1.99; and it was considered ‘No Identification’ when the value of the score was equal or lower than 1.69 **(29)**.

To perform data analysis, ClinProTools software (version 3.0, Bruker Daltonik GmbH, Bremen, Germany) and Flex Analysis v3.4 software were used. All spectra were exported as mzXML files using CompasXport CXP3.0.5 according to standard Bruker setup

### Data pre-processing

The following data pre-processing steps were carried out according to the literature **(30,31)**:

- baseline correction: a polynomial function fitted to a subsection of the baseline was used. For this, the “top-hat” option was selected, whose length was set at 10% of the minimum width of the baseline.
- spectra recalibration: recalibration was performed at 1,000 ppm maximum peak offset and 30% coincidence with the calibrating peaks, excluding null or out-of-range spectra.
- smoothing: with this function the noise level generated by the matrix and the components external to the sample was reduced in order to amplify the information contained in the spectra. Smoothing was performed in one cycle, on the order of 7 Da wide.

#### Unsupervised analysis models

-Principal Component Analysis (PCA): In order to evaluate the possible distributions or “clusters” on the isolates of both classes (PCV13 and NON PCV13), a PCA analysis was performed on the registered spectra. For this, the ClinPro Tools software (version 3.0, BrukerDaltonikGmbH, Bremen, Germany) was used. For this analysis, 10 principal components (CP) were defined for the entire spectral range (2000-20000Da).
-Hierarchical Cluster Analysis (HCA): From the generated spectra, a dendrogram was constructed based on the similarities of the peak intensity and mass signals. To carry out this point, the ClinPro Tools software (version 3.0, BrukerDaltonikGmbH, Bremen, Germany) was used.

#### Supervised analysis models

##### Peak Selection

To select the characteristic peaks of the two classes (PCV13 and NON PCV13) the following statistical tests were used: t-test/analysis of variance ANOVA (PTTA), Wilcoxon or Kruskal–Wallis test (W / KW) and Anderson test – Darling (AD). A P value of 0.05 was established as the cut-off point **(27)**:

-if p is <0.05 in the AD test, a characteristic peak is selected if the corresponding value of P in the W/KW test is also <0.05.
-if p is 0.05 in the AD test, then a characteristic peak is selected if the corresponding p value in ANOVA is also <0.05 **(32)**

Biomarker peaks were identified by class comparison using the “PeakStatisticTable” function in ClinPro Tools followed by manual confirmation, using Flex Analysis (Khot and Fisher, 2013) **(33).**

The discriminative power for each biomarker was further described by receptor operating characteristic (ROC) area under the curve (AUC) analysis. The ROC curve provides a graphical description of the specificity and sensitivity of a test, and in this case an assessment of the discrimination quality of a peak. An AUC value of 0 indicates that the considered peak does not discriminate, while an AUC of 1 indicates that the considered peak discriminates.

Three supervised classification models were calibrated using ClinPro Tools software: Genetic Algorithm (GA), Supervised Neural Networks (SNN) and Quick Classifier (QC). In turn, the QC algorithm was calculated using the ANOVA variance test (QC-ANOVA), the Kruskal-Wallis test (QC-WKW) and the Anderson-Wallis test (QC-DaV); all provided by ClinPro Tools software. For all cases, the selection of the maximum number of best peaks was set to 1000, and the maximum number of generations was set to 10. For the GA model, which selects the best peaks for classification, the k-NN (nearest neighbor) algorithm set to 3 was used for binary classification.

The developed calibration models were validated by cross validation in 10 iterations, leaving out 20% of the samples in each cross.

##### Independent validation of supervised classification models

To assess the robustness of the developed models, an independent set of isolates was selected for validation. A total of 100 isolates corresponding to the national surveillance of the 2020-2021 periods were used (Table S1).

##### Capsular typing

Gold standard of pneumococcal serotyping (the Neufeld-Quellung reaction) was performed using pool, group, type and factor specific commercial antisera produced by the Statens Serum Institute (Copenhagen, Denmark). Capsular types were assigned in accordance with the Danish nomenclature system **(8, 34,35)**

##### “Real time classification”

The strategy applied was based on the use of classification models in “real time”. Briefly, each *Streptococcus pneumoniae* isolate sent to the National Reference Laboratory for serotyping was previously challenged by the classifying models, and according to its prediction, either PCV13 or NON PCV13, the reference technique was carried out for capsular typing. The work algorithm can be visualized in figure 1.

**Figure 1.**
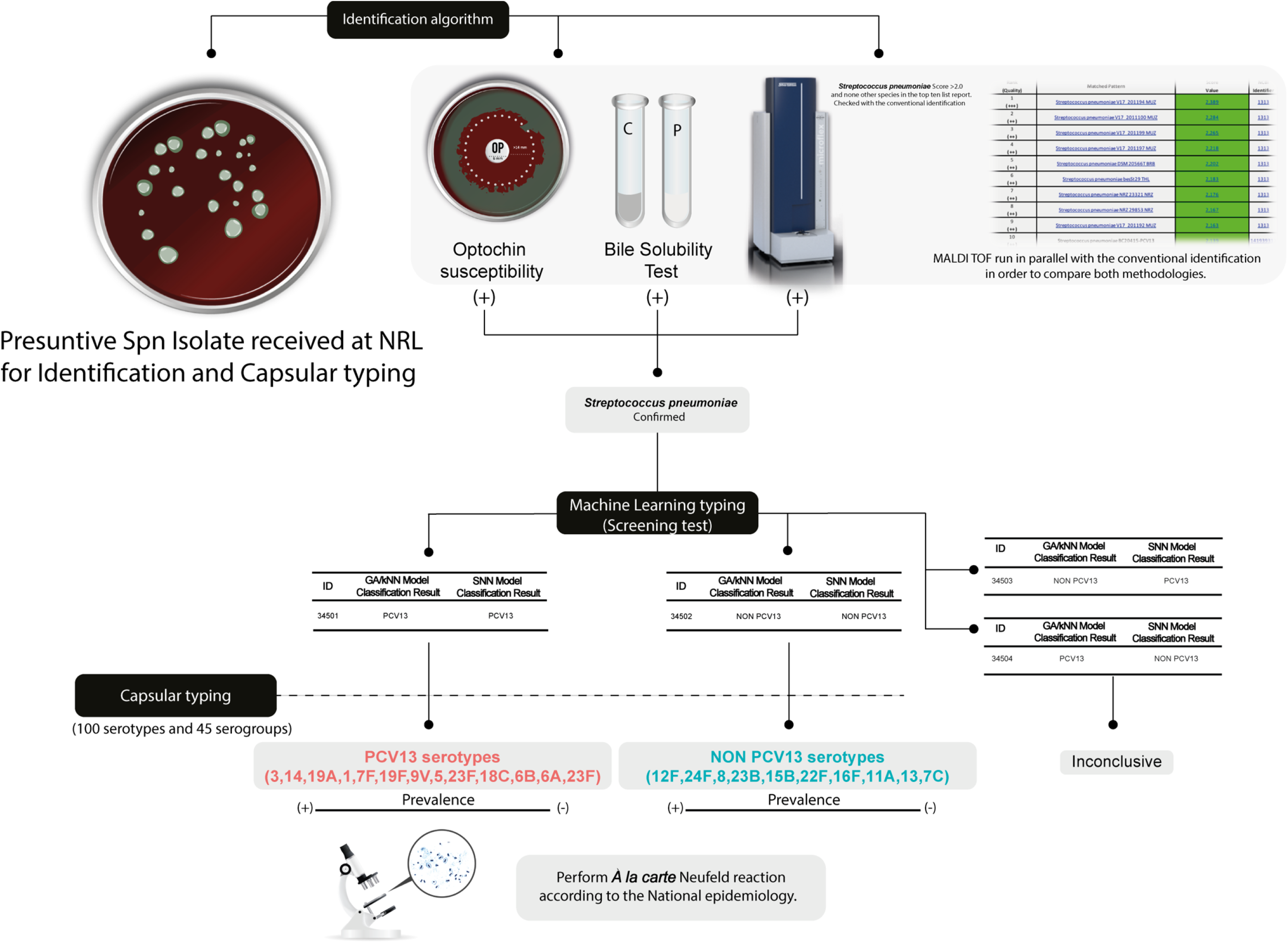
Workflow used for serotyping unknown isolates. In the first place, all isolate was processed by the conventional identification methods and MALDI-TOF (MicroFlex platform), if the results were in agreement, then, the spectrum was loaded into the ClinPro Tools software, in which, depending on the prediction of the models, we performed the personalized Quellung reaction according to the national epidemiology of circulating serotypes.

##### Statistical analysis

For statistical analysis, GraphPad Prism 9.0 software (GraphPad Software, Inc., San Diego, California, USA) was used. For the comparison of the classifying models with respect to the gold standard, contingency tables were made for each model (using binary variables), calculating sensitivity, specificity, precision, positive predictive value and negative predictive value. The Kappa index was calculated to establish the degree of concordance between the two methods. An index between 0.81 and 1.0 indicates perfect agreement, between 0.61 and 0.80 high or substantial agreement, between 0.41 and 0.60 moderate, between 0.21 and 0.40 slight, less than 0.20 insignificant **(36)**

In the same way, the CLSI guideline EP12-A2 **(37)** was used to compare methods that report results qualitatively. When the comparison is made with a method that is not considered a reference, the degree of similarity between the methods is measured through the percentage of negative agreement and the percentage of positive agreement. The diagnostic parameters of both methods are then compared to determine if the difference between the two is statistically significant.

## Results

### Unsupervised analysis

Principal Component Analysis (PCA) was performed for the 23 isolates used as training set. Figure 2 shows the graph of the accumulated variance for the new components. In this case, 10 principal components (PCs) were defined, of which the first three explained 82% of the variance of the spectral data. In this way it was possible to reduce all the information contained in the MALDI-TOF spectra in a few new variables. This means that if the spectrum is thought of as a multivariate system, each peak or signal as a function of the charge mass (m/z) represents a variable, then the PC’s represent new variables that contain all the information of the multivariate system (spectrum).

**Figure 2.**
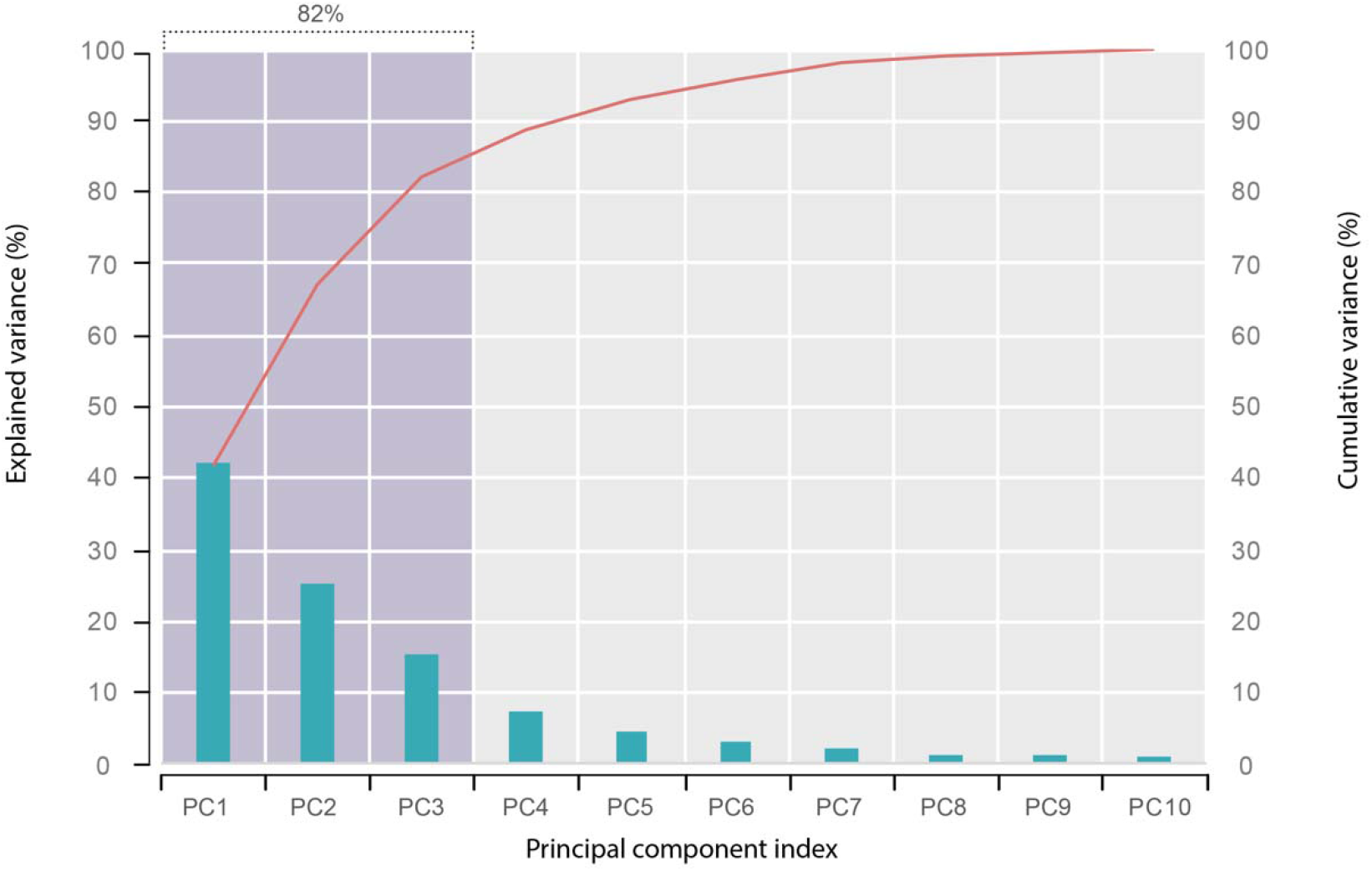
Variance attributable to each principal component generated and percentage of accumulated variance.

This allows you to graphically represent all the spectra together in three and two dimensions on the Score plot.

Figures 3 and 4 shows the score plots for the PC’s. In section A the first three components can be graphed simultaneously in a three-dimensional way, while in sections B, C and D the component 1, 2 and 3 can be seen combined in two dimensions.

**Figure 3.**
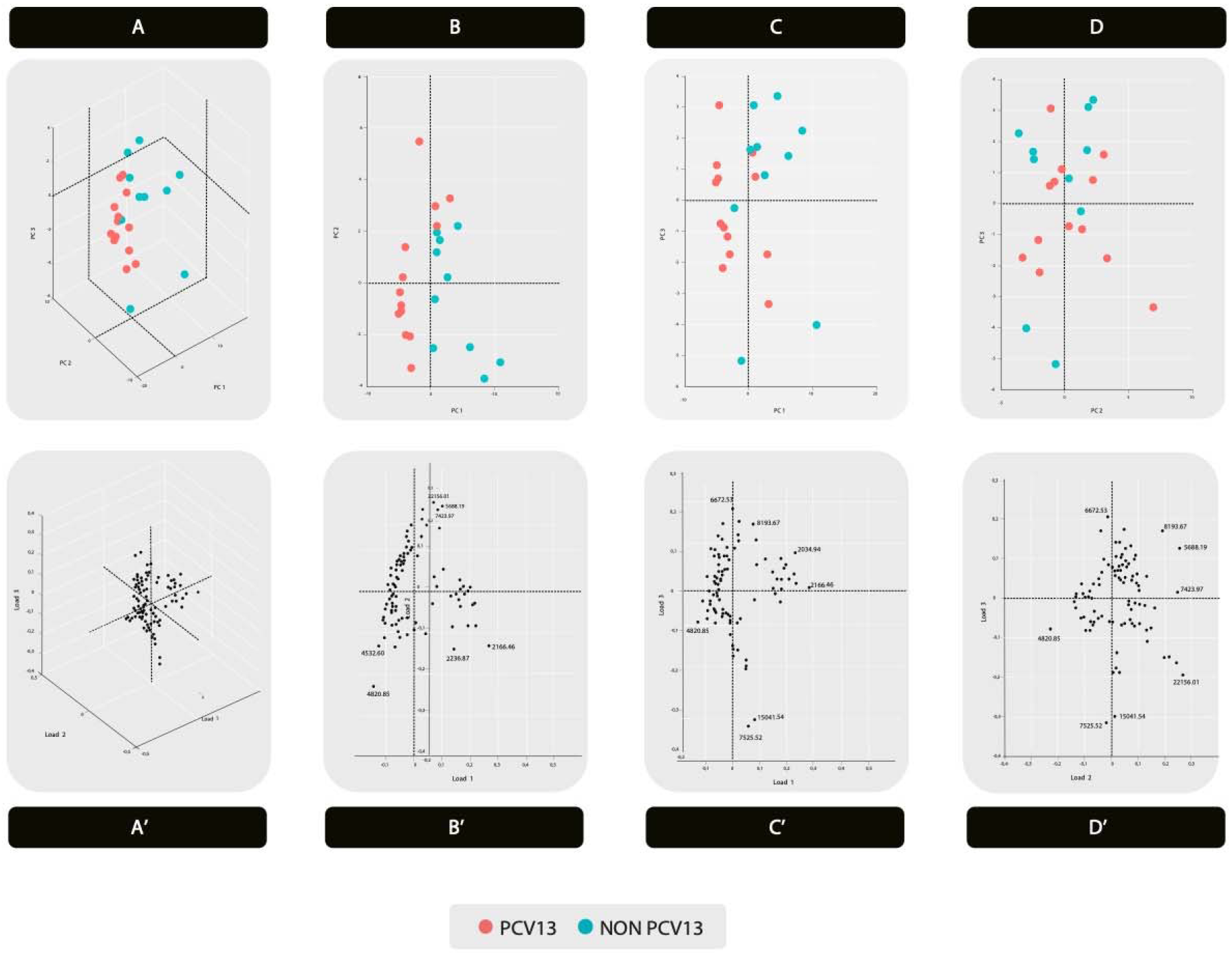
Score plot. Dimensional image of PCA showing the distinction between the 23 isolates; 13 PCV13 isolates (pink) and 10 NON PCV13 isolates (green). Representation of the loading plots of the PCs. The most significant variables are labeled with their corresponding m/z.

**Figure 4.**
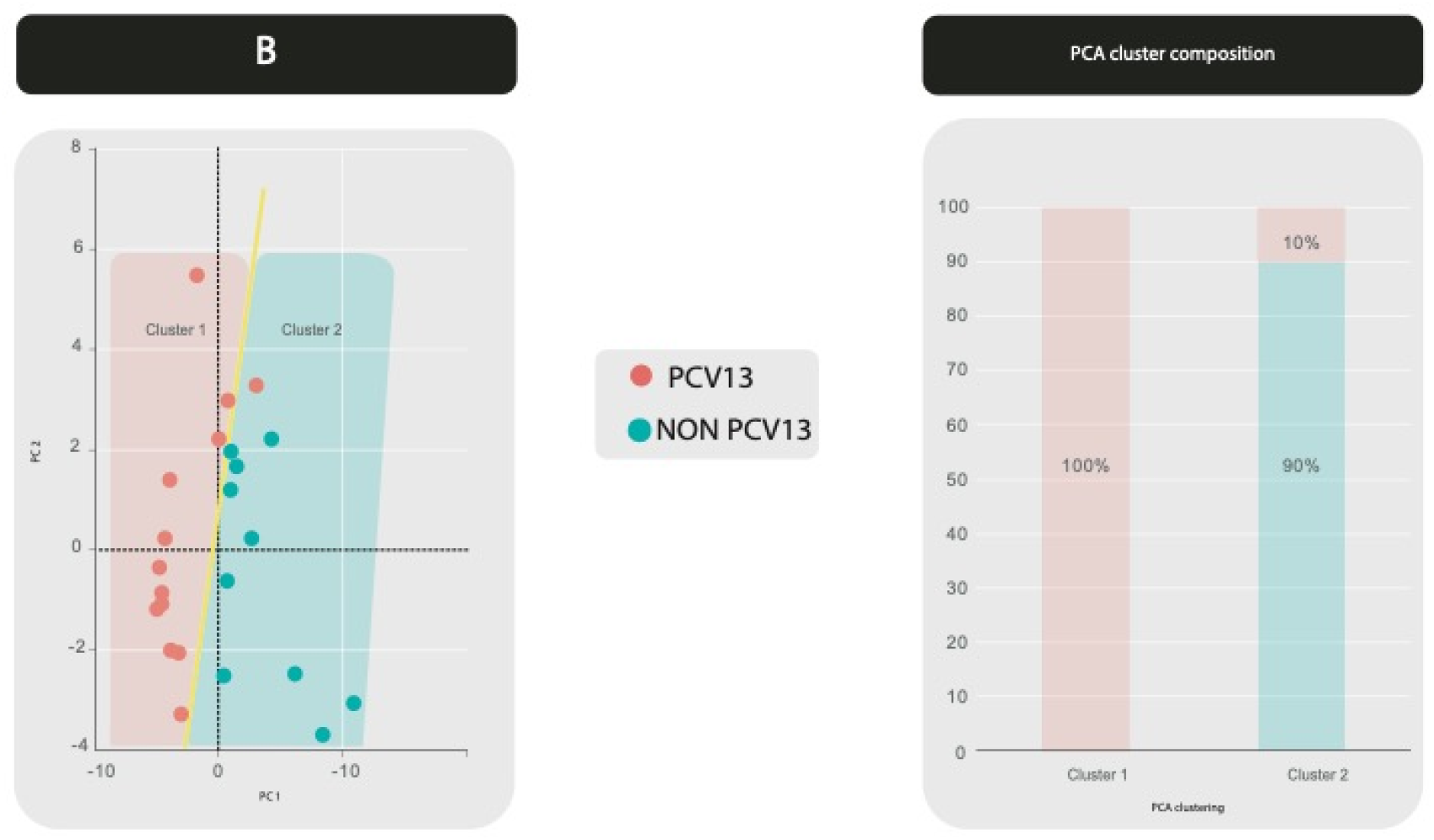
Unsupervised statistical analysis. Dimensional image of PC1 versus PC2 showing the distinction between the 23 isolates; 13 PCV13 isolates (pink) and 10 NON PCV13 isolates (green).

Clusters appeared as two slightly overlapping groups. Specifically, cluster 1 achieved 100% (12/12) of homogeneity for spectra corresponding to PCV13 serotypes and 92.3% (12/13) of PCV13 spectra fell in this cluster, while for cluster 2 a 90% (10/11) homogeneity was achieved for spectra corresponding to NON PCV13 serotypes group and 100% (10/10) of this group spectra fell in this cluster (Fig. 4). This result shows that both groups may have distinctive protein signatures that allow their discrimination.

At the same time, an unsupervised hierarchical clustering analysis (HCA) was carried out, from which the dendrogram that can be seen in Figure 5. The horizontal axis represents the distance calculated in the clustering, which is shown in relative units, corresponding to the similarity of the MALDI-TOF spectra.

**Figure 5.**
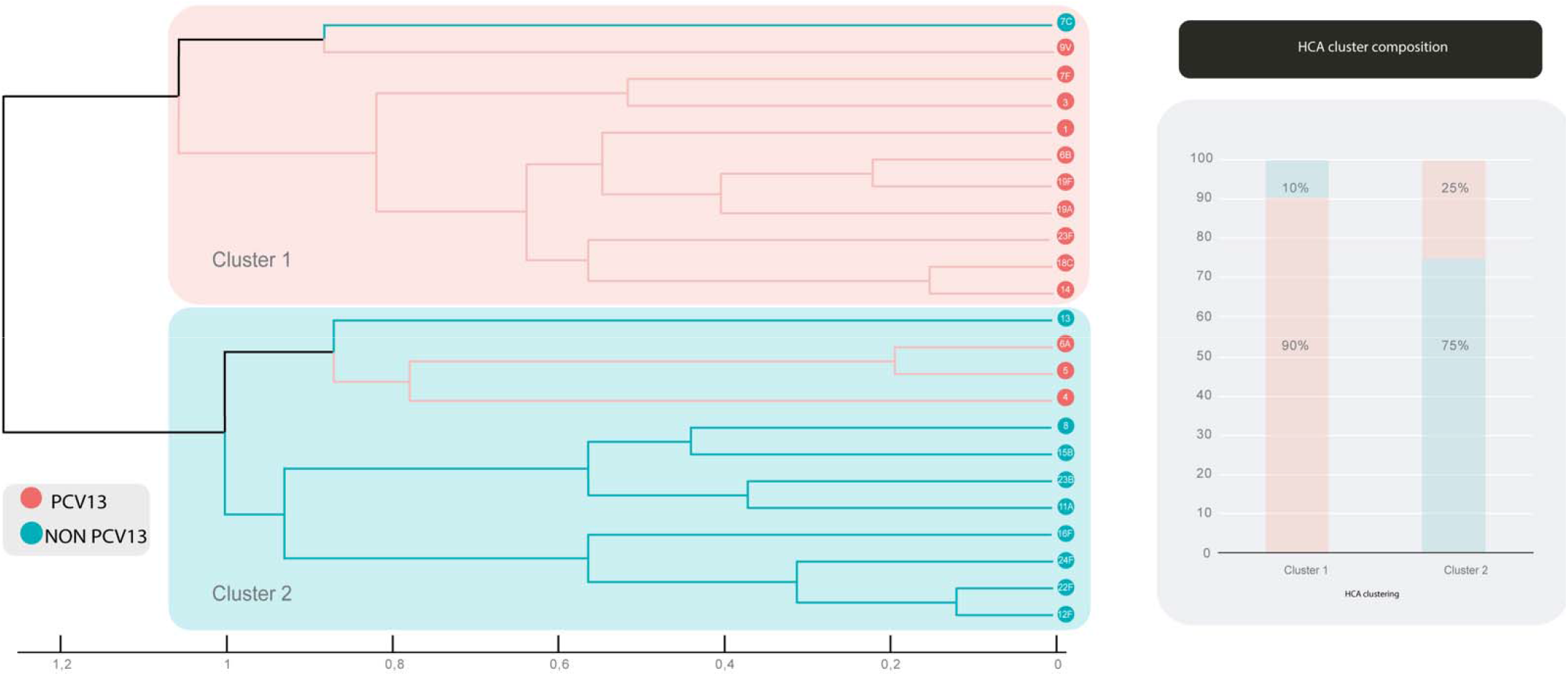
Dendrogram corresponding to unsupervised hierarchical clustering

Figure 5, shows the visualization of the respective relationship between the isolates. Based on a distance approximate above of 1.1, two clusters were present. Cluster 1 achieved 90% (10/11) of homogeneity for spectra corresponding to PCV13 serotypes and 77% (10/13) of PCV13 spectra fell in this cluster, while for cluster 2 a 75% (9/12) homogeneity was achieved for spectra corresponding to NON PCV13 serotypes group and 90% (9/10) of this group spectra fell in this cluster.

### Supervised analysis models

Subsequently, with the additional information of each isolate to define each class (PCV13/ NON PCV13), the supervised multivariate analysis was performed.

To recognize the spectral patterns of each class, all the recorded spectra were imported into the Clin ProTools software. Data were entered into two groups (PCV13 and NON PCV13) according to the results of previously obtained Quellung serotyping.

Figure 6-A shows the two-dimensional distribution graph of all the spectra of each class based on the two best peaks obtained for their classification; which were 170;10786 Da and 78;4703 Da (Fig.7). The peak number and m/z values of these are shown on the x and y axes, while the ellipses represent the 95% confidence interval. On the other hand, figure 6-B shows the ROC curves of the two selected peaks. The area under the curve (AUC) represents the discriminatory potential of each biomarker peak.

**Figure 6.**
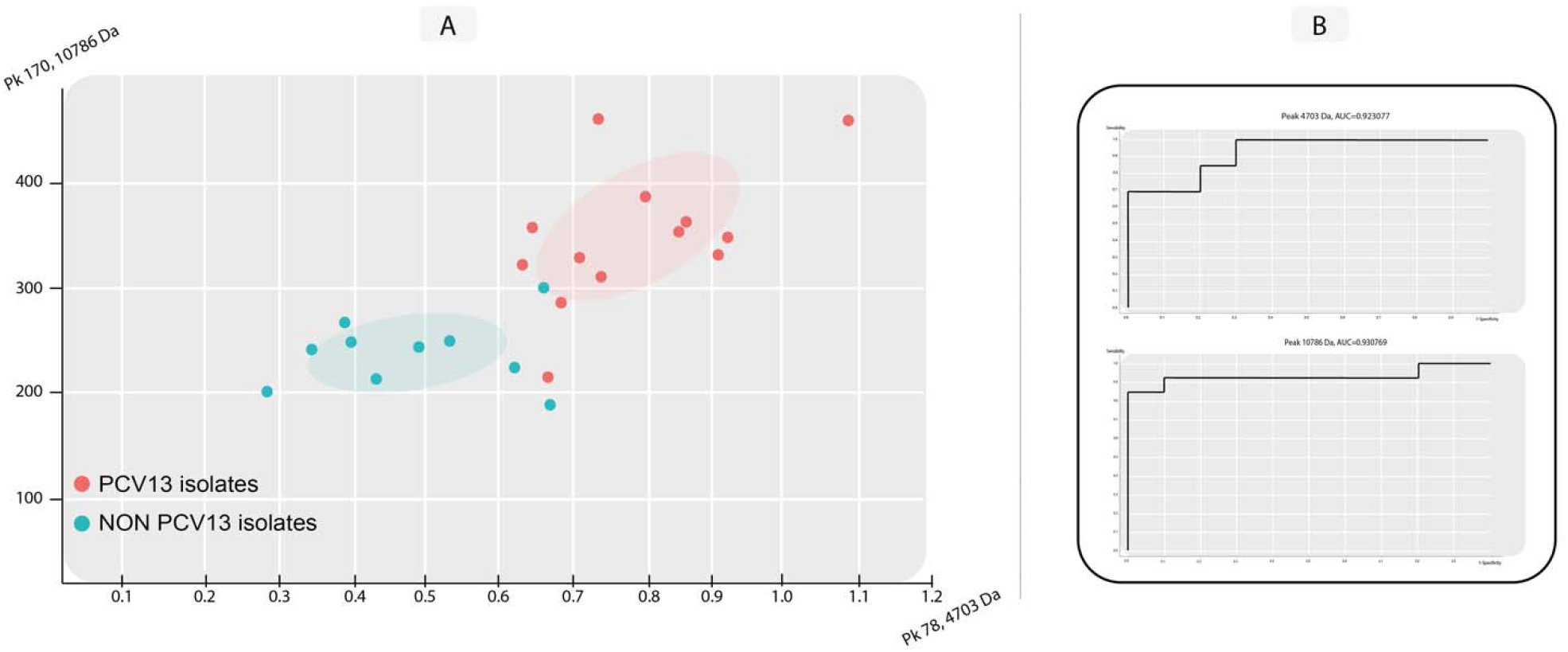
**A-** Two-dimensional distribution of the characteristic peaks of each generated class. **B-** ROC curves with their corresponding AUC values of those peaks.

**Figure 7.**
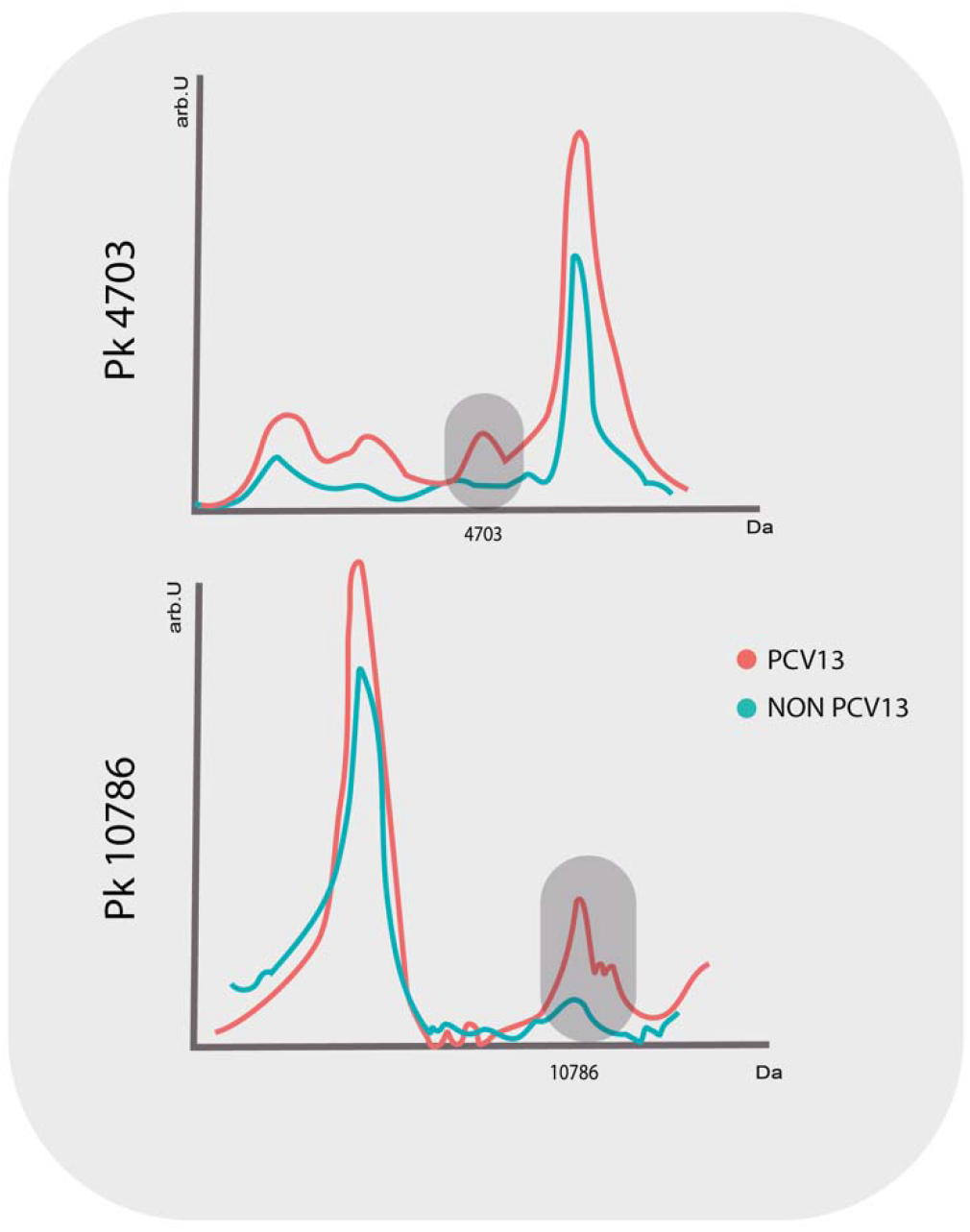
Graphical representation of the 2 best discriminatory peaks: Pk 4703 Da and Pk 10786 Da.

The best 10 peaks calculated by the software, with their statistical values, are summarized in Table 2.

**Table 2.** The best 10 peaks calculated by ClinPro Tools software.

| ***Index*** | **Mass** | **Dave** | **PTTA** | **PWKW** | **PAD** |
| --- | --- | --- | --- | --- | --- |
| 78 | 4703.69 | 0.57 | 0.00299 | 0.00302 | 0.807 |
| 170 | 10785.95 | 0.15 | 0.00299 | 0.00358 | 0.423 |
| 81 | 4794.85 | 0.57 | 0.00693 | 0.00302 | 0.194 |
| 180 | 1356.74 | 0.2 | 0.00739 | 0.00358 | 0.0895 |
| 116 | 6253.35 | 0.3 | 0.0121 | 0.0188 | 0.635 |
| 117 | 6270.43 | 2.56 | 0.0127 | 0.0211 | 0.121 |
| 43 | 3173.19 | 1.02 | 0.0168 | 0.0188 | 0.132 |
| 118 | 6436.73 | 0.65 | 0.0176 | 0.0188 | 0.875 |
| 171 | 10991.5 | 0.55 | 0.0176 | 0.0188 | 0.817 |
| 39 | 2983.78 | 1.15 | 0.0176 | 0.0188 | 0.192 |

DAve=Difference between the maximum and minimum intensity of the average peak of all classes;
PTTA= p-value obtained through the t-test, range 0-1; where 0: good and 1: bad
VPWKW= p value obtained using the Wilcoxon / Kruskal-Wallis test; range 0-1; where 0: good and 1: bad
PAD= p-value obtained by the Anderson-Darling test: range 0-1; 0: non-normal distribution, 1: normal distribution

In Figure 7, it can be seen that the two best peaks selected according to their statistical parameters, which were used by the software to perform Figure 6, are found in the PCV13 isolates and that they are practically absent in the NO PCV13 isolates.

### Classification Models

A total of 5 algorithms were calculated: GA/kNN, SNN, QC-DAv, QC-Anova and QC-WKW, the results of the different parameters of each algorithm are summarized in Table 3.

**Table 3.**
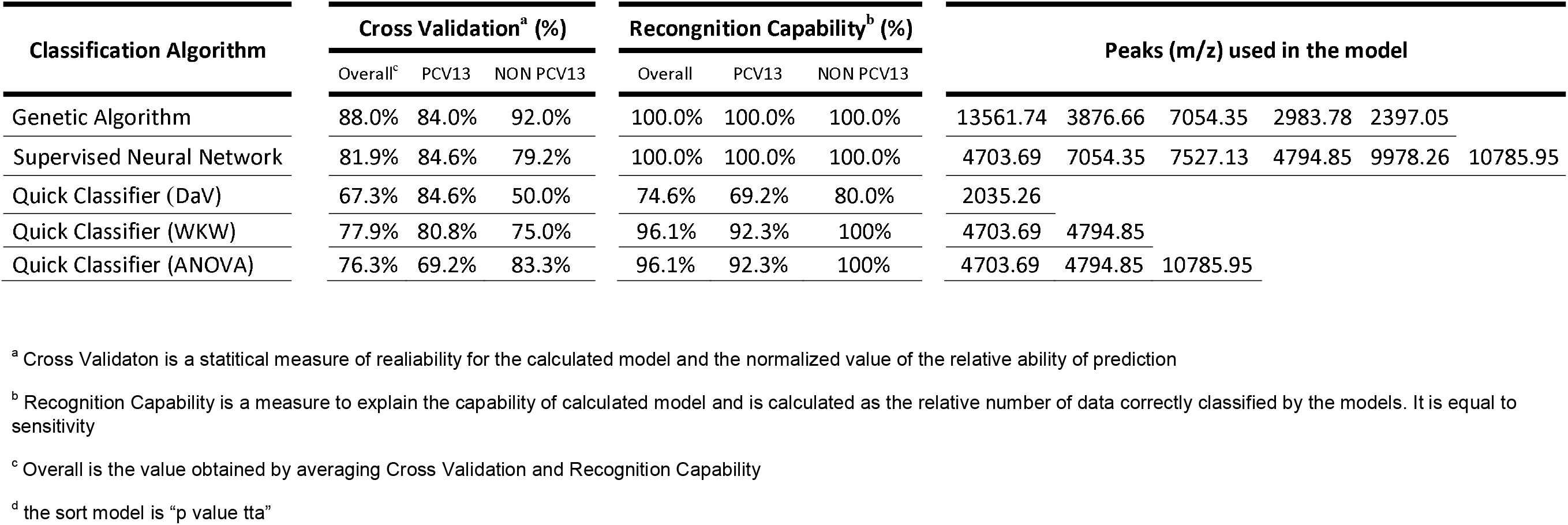
Parameters of each calculated model.

Based on the results obtained, it was decided to use only GA/kNN and SNN since they presented the highest values of theoretical recognition.

For the validation of the selected models, 100 isolates were used (Table S1) of which the serotype was unknown. The results of the performance of the predictive models with respect to the gold standard are summarized in Tables 5–7 and Figure 8.

**Figure 8.**
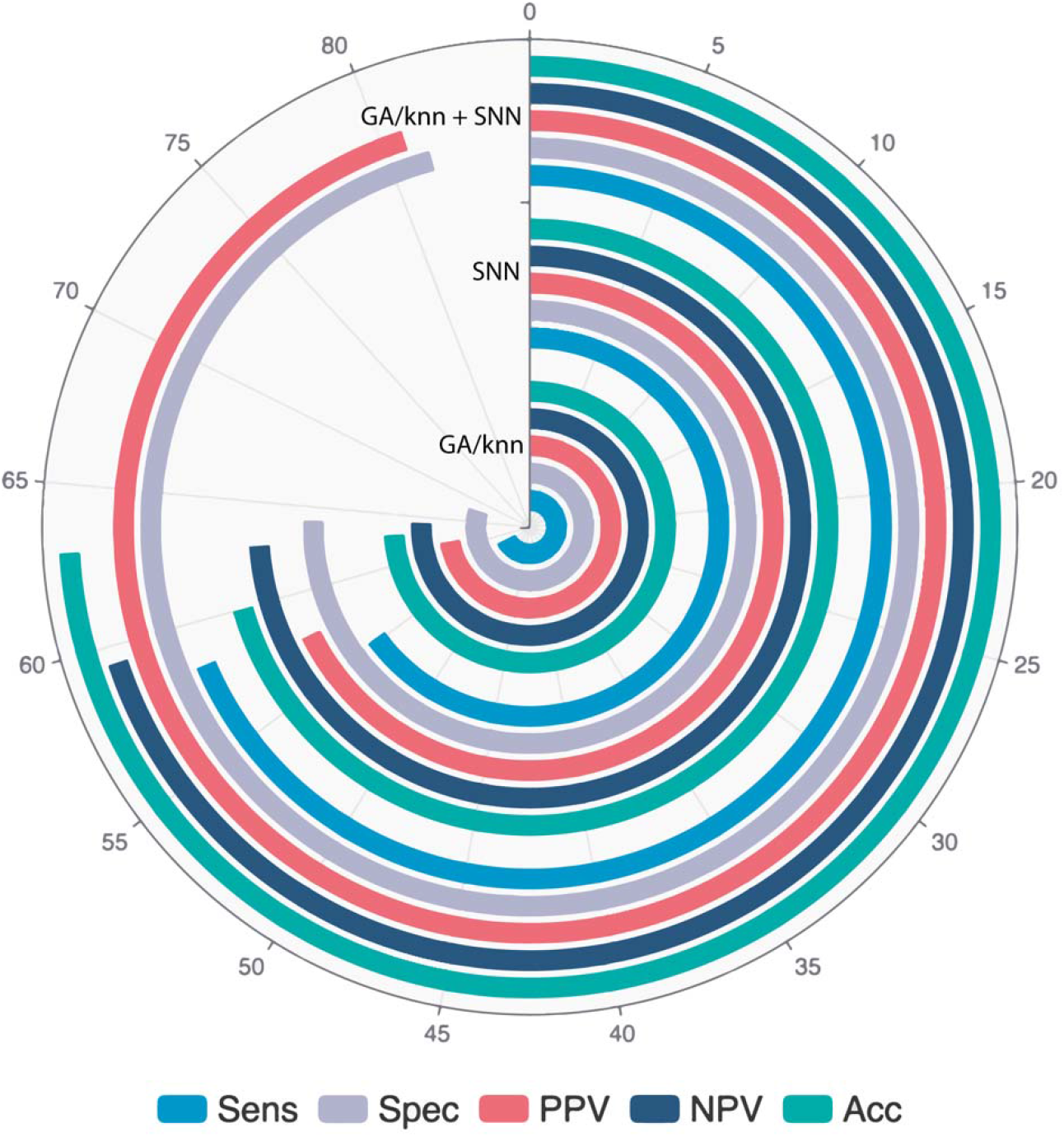
Radial representation of the different parameters evaluated.

**Table 5.**
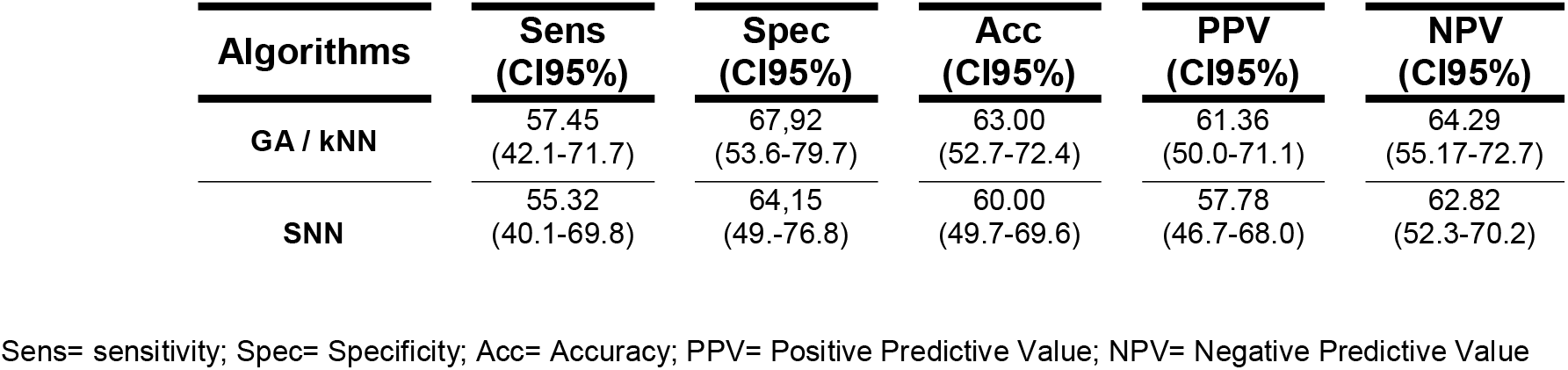
Results of the parameters evaluated for the selected algorithms.

**Table 6.** Results of the parameters evaluated considering only the concordances between both models. (n=63)

| **Algorithm** | **Sens**<br>**(CI95%)** | **Spec**<br>**(CI95%)** | **Acc**<br>**(CI95%)** | **PPV**<br>**(CI95%)** | **NPV**<br>**(CI95%)** |
| --- | --- | --- | --- | --- | --- |
| **GA / kNN** | 58.33<br>(40.7-71.7) | 81.48<br>(53.6-79.7) | 63.00<br>(61.9-93.7) | 80.77<br>(64.5-90.6) | 59.46<br>(48.9-69.2) |
| **SNN** |  |  |  |  |  |

**Table 7.**
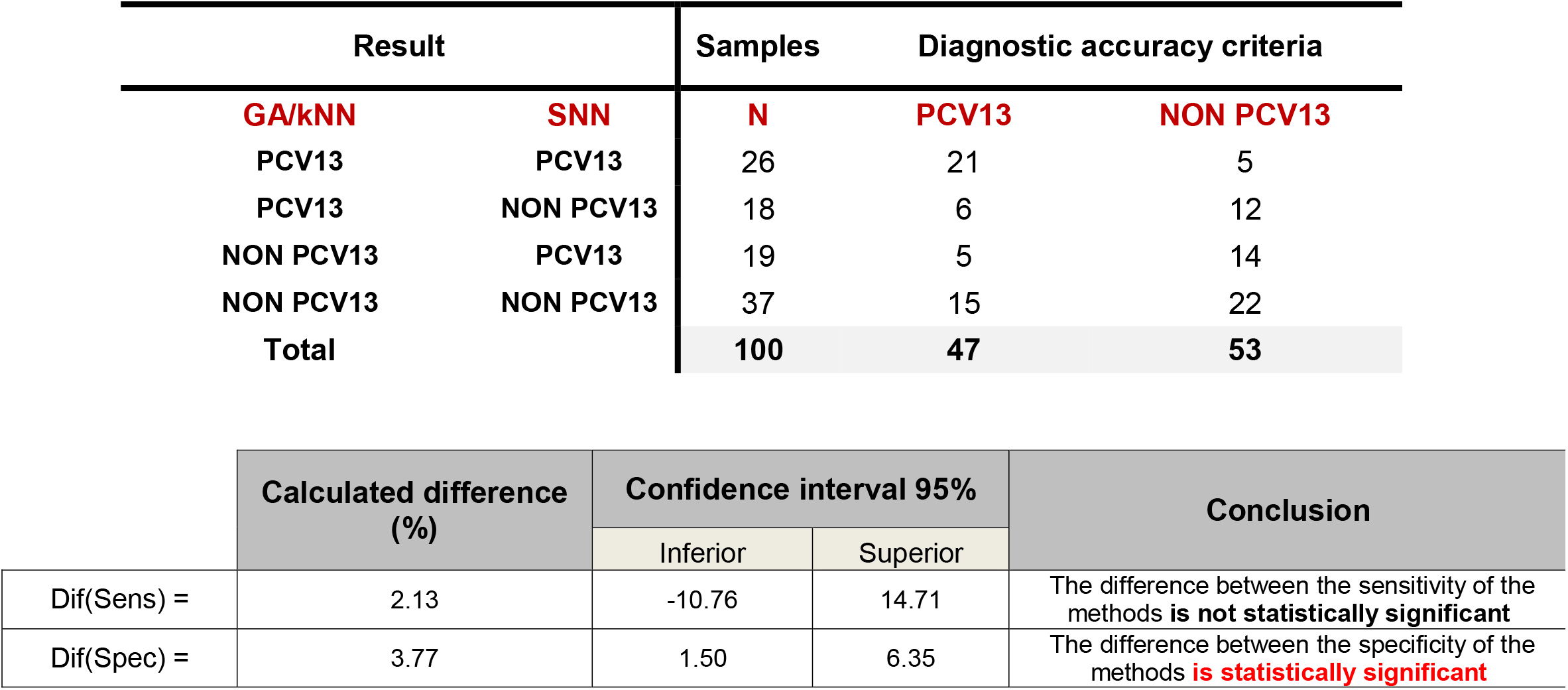
A Three-Way Comparison between the candidates methods and diagnostic accuracy criteria.

Table 6 shows the statistical parameters of the GA/kNN plus SNN algorithms with respect to the gold standard, considering only the isolates that showed concordance between both classifying models (63%).

It could be seen that 37% of the isolates could not be categorized, resulting inconclusive by this approach.

Table 7 shows the comparative results between the sensitivity and specificity parameters of the two predictive models with respect to the gold standard. It was observed that for the sensitivity parameter the differences were not statistically significant, while for the specificity the differences were significant.

Regarding the Kappa index, it was possible to determinate that the degree of concordance between the two predictive models was weak.

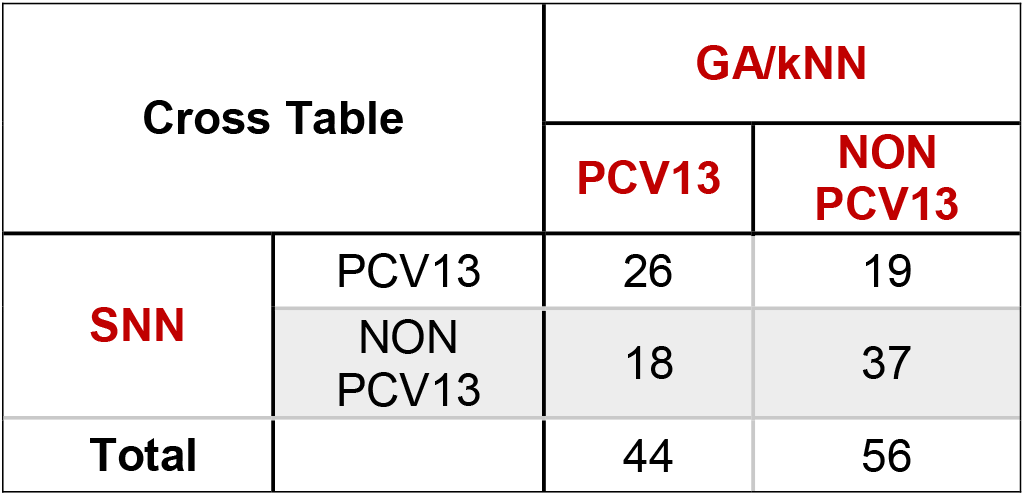
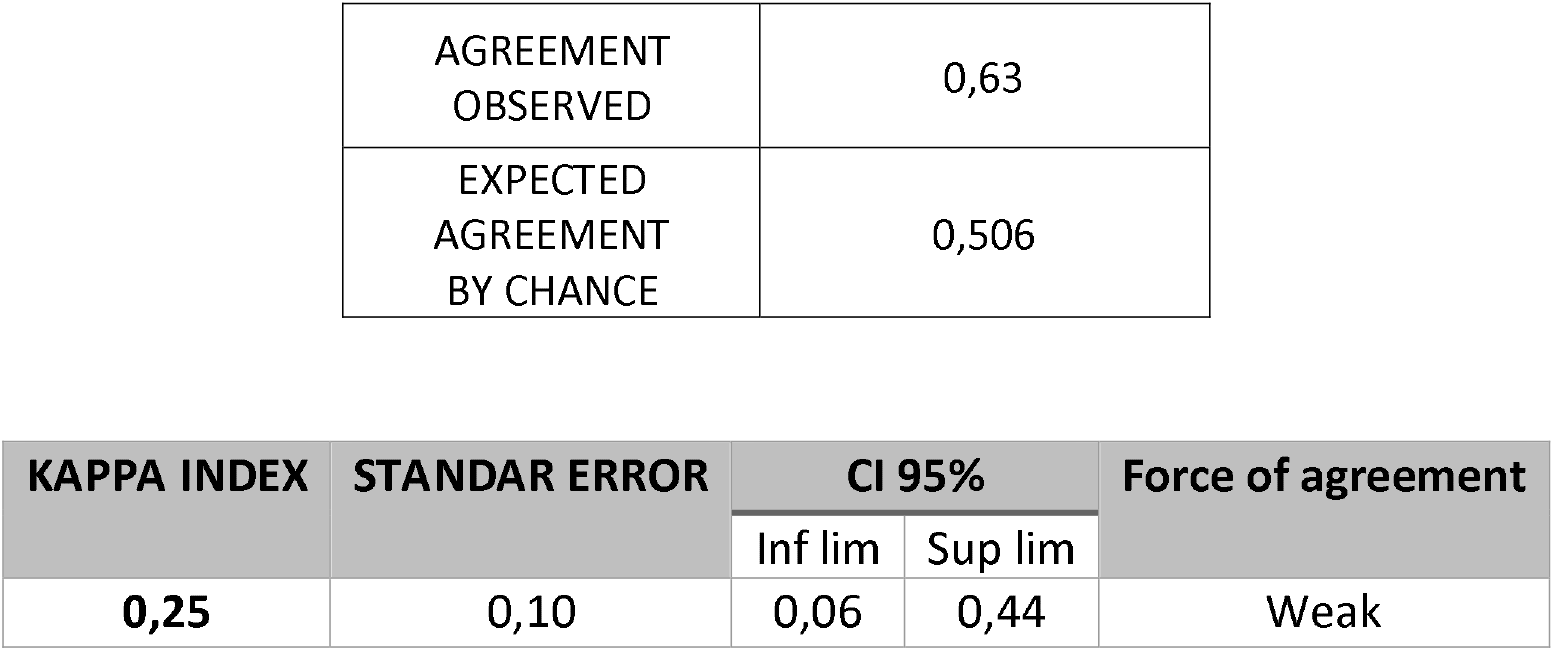

| Cross Table |  | GA/kNN |  |
| --- | --- | --- | --- |
|  |  | PCV13 | NON PCV13 |
| SNN | PCV13 | 26 | 19 |
|  | NON PCV13 | 18 | 37 |
| Total |  | 44 | 56 |

|  |  |
| --- | --- |
| AGREEMENT<br>OBSERVED | 0,63 |
| EXPECTED<br>AGREEMENT<br>BY CHANCE | 0,506 |

| KAPPA INDEX | STANDAR ERROR | CI 95% |  | Force of agreement |
| --- | --- | --- | --- | --- |
|  |  | Inf lim | Sup lim |  |
| 0,25 | 0,10 | 0,06 | 0,44 | Weak |

## Discussion

Invasive disease caused by *Streptococcus pneumoniae* (pneumococcus) is a serious infection. It can produce a wide spectrum of clinical manifestations including sepsis, meningitis, bacteremic pneumonia, arthritis, osteomyelitis, cellulitis, and endocarditis.

The people with the highest risk of suffering from this pathology are those under 2 years of age, the elderly and people with immune disorders or certain respiratory, cardiac, renal pathologies, among others.

Pneumococcal vaccination is intended to reduce the incidence, complications, sequelae, and mortality from pneumonia and invasive pneumococcal disease. Two types of vaccines are available: a polysaccharide vaccine against 23 serotypes (PPSV23) and a conjugate vaccine against 13 serotypes (PCV13) **(13).**

Currently, serotype identification is performed by the Quellung technique. This technique requires expensive reagents, is very laborious, requires specialized staff and involves long times until the result is obtained. In Argentina the determination of the capsular type is performed only in the NRL and, due to this, it is difficult to obtain the results in real time. For all the above reasons, the search for reliable, fast and cheap alternative methods for Spn serotyping is of great interest in the field of public health.

In this sense, in recent years MALDI-TOF mass spectrometry has revolutionized the field of clinical microbiology **(38)**. Although its application is fundamentally related to microbiological identification **(39)** applications combined with other bioinformatic analysis platforms have recently emerged **(40,41)**. In line with the above, the use of artificial intelligence has disruptively entered the field of health, thus becoming a new tool to be considered for the diagnosis of different pathologies **(42–44)**. The first attempts to discriminate Streptococcus from the viridans group, using mass spectrometry, were encouraging according to the published results which were based on a small number of isolates or creating a specific database for this group of microorganisms (**45,46)**. However, it was later found that the Bruker Biotyper 3.0 system could not resolve the low specificity in the identification of this genus **(47–50)** and why several authors have ventured into various approaches to overcome this limitation. In 2012, Werno et al **(51)** proposed the use of specific peak analysis to confirm identification, which implied an improvement in the typing of different species of *Streptococcus mitis*, including *Streptococcus pneumoniae*.

Subsequently, Ikryannikova **(52)** refers to the difficulty of discriminating pneumococcus from *Streptococcus mitis* by mass spectrometry; for which the authors proposed to use, in addition to biomarker peaks, artificial intelligence classifier algorithms. However, other authors **(53–55)** tried to replicate this methodology but were unable to find the peaks described.

It is important to note that this limitation occasionally arose during the creation of the spec-tral database and during the “real-time” classification of the unknown isolates performed in this manuscript.

In this context, an inclusion criterion was established for the isolates to be incorporated in this work (either as part of the training set as well as in the validation set). In this way, only those isolates that did not present discrepancies in the top ten of the results obtained by MS and that also showed a score > 2.0 were included, by doing this, we could affirm that our spectra accomplished the quality we needed to perform this approach.

The objective of this work was to evaluate the application of MALDI-TOF MS in combination with artificial intelligence algorithms, as screening methods in the serotyping of *Streptococcus pneumoniae*

To this end, two-class models, developed by machine learning algorithms, were proposed to differentiate PCV13 vaccine serotypes from NON PCV13 serotypes, which would allow to performed the Quellung serotyping in a more targeted manner combined with the most prevalent circulating serotypes in Argentina, substantially saving both reagents and man-hours.

First, a calibration set comprised of isolates of all the most frequent vaccine and nonvaccine serotypes within the local epidemiology was used, and a principal component analysis (PCA) was performed from the spectra obtained by MS. This unsupervised analysis was carried out with the aim of exploring the behavior of these objects (Spn isolates) in function of the new variables defined by this study. It was possible to see that the first three components explained the highest percentage of the variance of the data, which gave an accumulated variance of 82%.

In the PC1 vs PC2 score graph **(Fig. 4)**, a homogeneous distribution along the PC1 axis can be observed. This trend allowed us to discriminate two clusters that can be correlated with vaccine and non-vaccine isolates.

According to the loadings graph, the load values of each peak were obtained during the calculation of the main components. In this way, the peaks that give the greatest weight to the main components can be associated with the distribution of the samples observed in the score graph. This tool is useful for explaining the outliers that score charts show. From this it is possible to recalculate the analysis incorporating or leaving out the influential peaks according to the objective that is set. This analysis allows us to make decisions regarding which variables can be given or removed more weight to obtain the desired distribution.

In the unsupervised hierarchical clustering analysis **(Fig. 5)**, the spectra were compared with each other in pairs, and the value obtained from said comparison allowed the hierarchical clustering in branches within a taxonomic tree according to the proximity between them. In the dendrogram obtained, a high percentage of discrimination between both classes can be seen.

Once the behavior of the study objects was explored in an unsupervised manner, supervised training was implemented, which aims to give a predictive approach to analysis, considered a subdomain of artificial intelligence, in which the computer uses algorithms to learn from a set of past data to make predictions about new data. In this way, it is possible to classify according to a previously established criterion in the training or calibration phase.

First, the best discriminatory peaks for each class were identified, according to the previously established parameters, which yielded two biomarker peaks for each class (4703.69 Da and 10785.95 Da). The performance values observed in the detection of these peaks were acceptable, since two well-defined groupings were observed in the two-dimensional plot of classes **(Fig. 6)**.

Nakano et al **(56)** used ClinPro Tools software to create prediction models for the ten most prevalent serotypes in Japan, but were only able to validate the assay for three serotypes (3, 15A, and 19A). Subsequently Pinto **(57)**, through the use of biomarker peaks and using the BioNumerics 7.6 software, found encouraging results but only with the serotypes 6A, 6B, 6C, 9N, 9V and 14. However, both authors conclude that it is necessary to carry out an external validation to corroborate the true reproducibility of the evaluated approaches.

In this work, five classifier models were calibrated using the ClinPro Tools software, of which the recognition capacity and cross-validation values showed greater efficiency for the GA/k-NN and SNN algorithms. An important result to highlight is the following, and it is that in accordance with what was found in the unsupervised analysis, more precisely in the two-dimensional figure, it is that the peaks selected to be able to make this figure, are in agreement with the peaks used by some of the classifying models, providing more robustness to the results obtained.

When implementing these models independently, low sensitivity and specificity values were observed, which is an inappropriate option to use them in this way as a screening technique for the serotyping of unknown isolates. However, when applying both algorithms in parallel and combined, a notable improvement in specificity (82%) and therefore in the positive predictive value (81%) was achieved. However, negative sensitivity and predictive value values continued to be around 60%, yielding inconclusive results in 37/100 isolates.

As a perspective to improve the predictive parameters, we can mention the increase in the number of isolates both to create the training set as well as the number of isolates to challenge.

Although the results obtained for this thesis work were not as expected, by implementing the models developed in the form of screening, the use of antisera was reduced by 10.2% compared to the blindly Quellung technique.

Undoubtedly, the development and application of EM has contributed to meeting the demands in the field of microbiological diagnosis due to the important advantages it presents compared to more traditional methodologies. Among these advantages, its versatility can be mentioned (since it can be applied to bacterial, mycological, virus, parasite cultures, even from the clinical sample itself). Additionally, minimal sample preparation is required and it is a fast, easy-to-use analysis technique that allows parameters to be evaluated in real time **(58)**.

In this work it was possible to demonstrate that the combination of MALDI-TOF mass spectrometry and multivariate analysis allows the development of new strategies for the identification and characterization of Spn isolates of clinical importance. MALDI-TOF mass spectra generate a large amount of data, which requires appropriate analysis methods to make the most of the information contained in them. In this sense, multivariate analysis models (both supervised and unsupervised) allow extracting information from the spectra (multivariate data set) and correlating it with different properties of the samples.

The results of this work represent the bases to continue exploring the combination of MALDI-TOF mass spectrometry with multivariate analysis, with prospects of improving the predictive parameters so that the developed models are robust and reliable. On the other hand, the development of this thesis provided a solid background in multivariate analysis as a tool to extract useful information and produce inferences from a large amount of data.

Finally, the possibility of projecting these methodologies to the study of other pathogens constitutes an added value as a future perspective.

